# Microbial Remediation of Pesticide Accumulation and its Response of Bacterial Wilt in Brinjal

**DOI:** 10.1101/2023.05.22.541768

**Authors:** Shenaz Sultana Ahmed, Popy Bora

## Abstract

Growing brinjal (*Solanum melongena* L.) in a soil polluted with organophosphorus pesticide (OP) residues coupled with an additional threat of wilt disease caused by *Ralstonia solanacearum* (*Rs*), pose a formidable management challenge for residue free crop production. Our study aimed at identifying efficient OP-degrading bacteria (OPDB) and their compatibility with microbial bio-control agents (MBCA) for twin objective of microbial degradation of OP -residues and reduction in incidence of bacterial wilt of brinjal in OP-contaminated soil. As many, ten bacterial isolates showing OP-degrading potential were recovered through colony growth in mineral salt (MS) - medium treated with 25 ppm chlorpyriphos. Efficient isolates displaying growth up to 700ppm chlorpyriphos were further screened for OP-degradation, leading to identification of two most efficient OPDB, *Acromobacter marplatensis* [MW397524] (*Am*) and *Pseudomonas azotoformans* [MW397525] (*Pa*). These two isolates having *in vitro* compatibility with each other, showed further compatibility with two most widely used MBCA, viz., *Pseudomonas flourescens* [KT258013] (*Pf*) and *Trichoderma harzianum* [ON364138] (*Th*), facilitating the development of consortium having extended functional corridor. The response of combination of OPDB + MBCA(*Am* +*Pa* + *Pf* + *Th*) showed 80% lower percent wilt incidence (PWI), 2.8 times higher fruit yield of brinjal, and 15-25% lower OP-residues over un-inoculated control treatment. Our studies, hence, put forward a strong unified delivery mechanism of OPDB and MBCA as a part of green technology for chemical residue -free vegetable production in contaminated soils.

**IMPORTANCE:** Microbes isolated from crop rhizospheres are known to exhibit multiple functions (plant growth promotion, antagonism against soil borne pathogens e.g., *Rs,* and acaricidal properties) depending upon type of stimuli by regulating the plant defense. Considering the limited past efforts on twin objective of cleaning up the pesticide residues accumulated in the soil (microbial removal of pesticide residues) and providing an additional safeguard against soil borne pathogen causing bacterial wilt in brinjal to collectively develop a microbial consortia mediated crop production system, devoid of any chemical residues. Such an effort adds strength to organic production system on pesticide contaminated soils. In the present study, we succeeded in identifying two most effective chloropyrifos residues degrading OPDB having compatibility two MBCA for developing consortia formulation, which aided in developing pesticide residue free soil, neutralising bacterial wilt pathogen (*Rs*), and creating a better nutrient supply for a well deserved production system. Such efforts would pave the way for developing more robust microbial formulations (with emphasis on entomopathogenic application) having dynamic microbial functions to replace any futuristic use of agrochemcials.

---

The success of agriculture in providing the food security is widely acknowledged in favor of high yielding varieties coupled with extensive use of xenobiotics, mostly the pesticides (1) as a part of full proof crop protection strategy. Use of pesticide, hence, has become inevitable in modern agriculture in order to feed the consistently growing population from almost exhausted arable land and water resources to meet an additional food demand (2).

Amongst different groups of chemical pesticides, organophosphorus pesticides (OP) as esters derived from phosphoric acids, are the most commercially favoredclass of pesticides, occupying 38% of world market (3). Initially, OP were designed to be used as chemical warfare agents during World War II, but soon made massive in-roads in agricultural fields (4). The chlorpyriphos, an extensively used OP is considered having residues moderately persistent in soil, even after months of application (5), despite their water soluble nature and hence, served as potential source of hazard to soil and water with varied genotoxic effects (5-7), comprising 385 million annual cases of unintentional pesticide poisoning worldwide (8). These figures warrant an alarm towards the accumulation of OP-residues (9-11). Microbes-mediated biodegradation of pesticide residues has received much greater attention in recent past, considering this process, an economically more viable than other alternative processes, involving chemical or physical degradation (10, 11). In microbial degradation of pesticides, the participating microbes utilise carbon (C) and/or phosphorus (P) released from degradation of OP (11).

The OP are ubiquitously used in field and horticultural crops against a wide range of insect pests viz., leafhopper, cutworm, grub, bugs, aphids, lepidopteran foliage feeders etc., apart from weeds (3,12). Amongst the vegetables, brinjal crop (*Solanum melongena* L.) receives a significant proportion of chemical pesticides, of which, OP are mostly used against various insect pests, more particularly against brinjal fruit and shoot borers (*Leucinodes orbonalis*) (13). In addition to these insect pests, bacterial wilt disease in brinjal caused by *Rs* (formerly known as *Pseudomonas solanacearum* E.F. Smith) is the major cause of concern imparting severe plant mortality coupled with heavy yield loss reported worldwide (14-16) due to widespread occurrence of disease, prolonged survival in soil, and unusually diversified host range (17), causing staggering economic loss to the tune of INR 500 crores (62 million US dollars) annually (18). Several methods used to control bacterial wilt of brinjal include: soil disinfection (19), soil amendment (20), and chemical control (21), however only with a limited successon account of low host resistance (22) considering *Rs* as soil-borne pathogen (23).

Previous studies have reported an effective control of wilt disease through application of MBCA like *Pf*, *Bacillus subtilis*, *B. amyloliquefaciens*, and *Th* against variety of soil-borne diseases, inclusive of bacterial wilt (24-27), involving mechanisms *viz*., production of antibiotics, cell wall degrading enzymes, siderophore-mediated suppression, production of toxic metabolites, over-crowding the pathogen, antibiosis, and induction of host resistance (28, 29) with a strong possibility of plant growth promotion ability as well. However, application of OPDB and MBCA lacks wide scale field application, majority of which are laboratory-based studies only. Hence, a suitable cost effective delivery mechanism of OPDB for contaminated soils and managing wilt disease of brinjal through MBCA, need an urgent attention for developing a double layer remediation strategy. The studies on these lines are miniscule in number. In this background, we carried out the studies with objectives comprising,, i. isolation and characterization of OPDB; ii. *in vitro* evaluation of OPDB against OP-degradation, iii. *in vitro* evaluation of MBCA against Rs, the wilt disease pathogen, and iv evaluation of different combinations of OPDB and MBCA against wilt incidence, changes in OP-residues, and agronomic performance of brinjal.

## RESULTS

### Bacterial wilt pathogen and pathogenicity manifestations

Pure culture of bacterial wilt pathogen (*Rs*) maintained in TTC-medium at 27±2°C (Fig. 1a) showed predominantly rod shaped cells (Fig. 1b). The pathogenicity of *Rs* under water culture and soil culture further demonstrated an epinasty growth pattern of inoculated leaves (downward bending of leaves) as first recognizable symptom, followed by wilting of first and second leaf from the apex leaf within three-to-four days of inoculation. While, wilting of lower-most leaves was observed as the last stage of development of wilt disease symptoms. The symptoms progressed, until the entire plant collapsed within 30-days of inoculation (Fig. 1c). Vascular discoloration coupled with longitudinal splitting of stem was observed in the internal tissues, with oozed out milky white bacterial mass in cut section, when placed in a test-tube containing clean water. Reisolated pathogen cultured in *Rs-*specific TTC-medium showing smooth, concave, mucoid, pink colonies was identified as *Rs* (Supplementary Table 1).

**Fig. 1.**
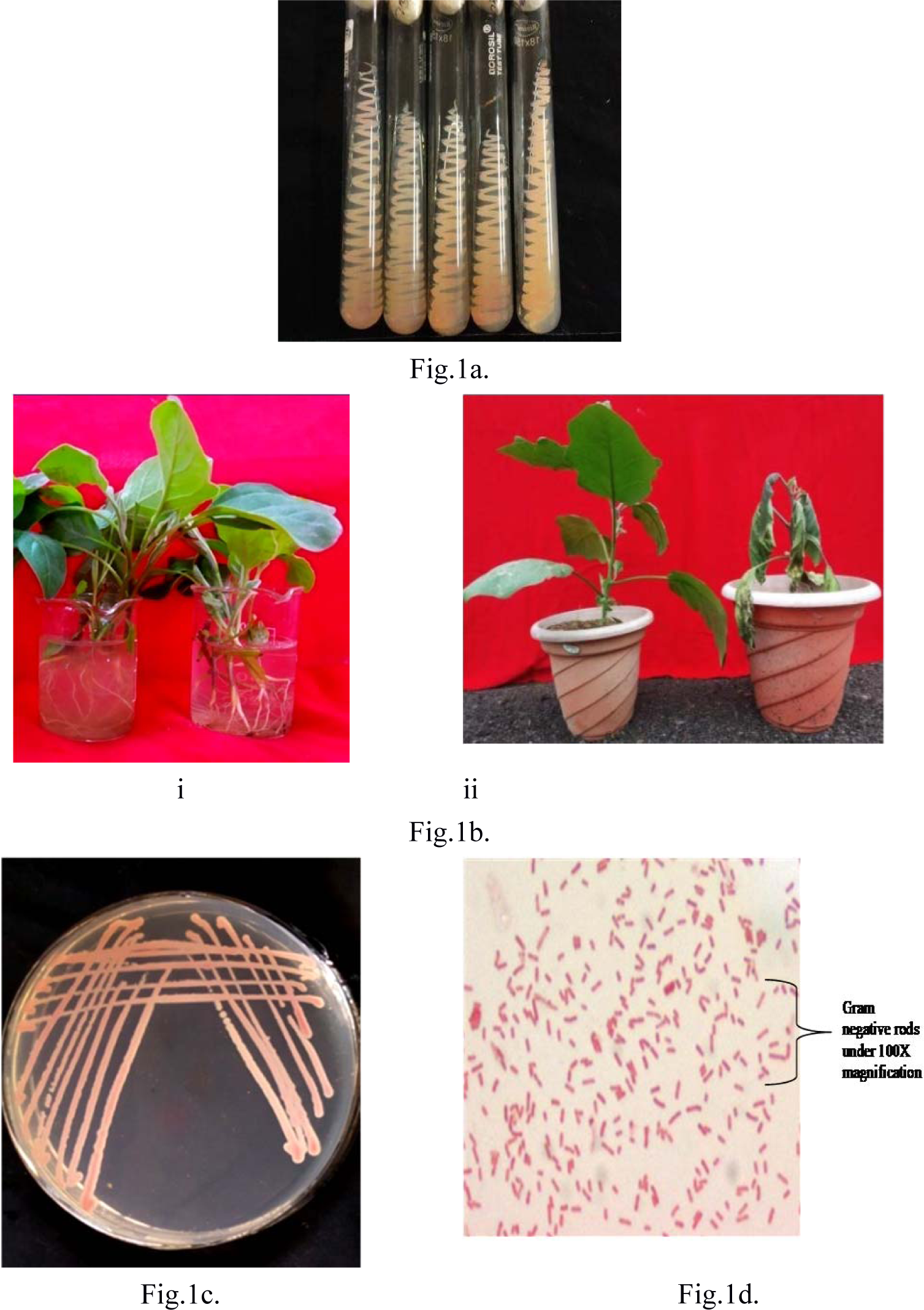
Reisolation and characterization of pathogen causing wilt disease in brinjal. Fig.1a. Pure culture of *Ralstonia solanacearum* (*Rs*) collected from Department of Plant Pathology, AAU, Jorhat, Assam (India). Fig.1b. Brinjal seedlings (one-month-old) treated with sterilized distilled water (i) and result of pathogenicity test (ii). Fig.1c. Pure culture of reisolated *Ralstonia solanacearum* (*Rs*) pathogen from inoculated brinjal seedlings. Fig.1d. Gram negative cells of reisolated *Rs*(100X) characterized by their pink coloured rod shaped bacilliform structure

### Identification and screening for OP-degradation

A total often different bacterial isolates were observed growing prominently in the soil samples collected from different brinjal fields based on their different colony morphology (Supplementary Table 2). These isolates were observed utilising OP-potential OPDB isolates were assigned with unique identification codes based on origin of soil samples *viz.*, TZ1, TZ2, TZ3 (Tezpur); JHT1, JHT2, JHT3, ICR1, ICR2 (Jorhat); and MG1, MG2 (Mangaldoi). Out of these 10 bacterial isolates, three isolates *viz*., JHT1, MG2 and TZ2 showed potential growth upto 700ppm of chlorpyriphos in MS medium (Fig. 3), while other two isolates viz., JHT2 and MG1 grew well upto 500 ppm of chlorpyriphos (Fig. 3). isolates (JHT1 and MG2) continued showing growth upto 700ppm. The growth kinetics of five efficient OPDB isolates (TZ2, JHT1, JHT2, MG1, and MG2) utilizing pesticide at 500 ppm revealed two isolates, JHT1 and MG2 showing highest growth (Fig. 4) and degraded 90-95% chlopyriphos (Fig. 5a-5d) within 45-days of inculation. These observations were further validated through analysis of residual chlorpyriphos and degradation metabolites at 0,3,5,7, and 14-days intervals following inoculation. Two isolates (JHT1 and MG2) hydrolyzed chlorpyrifos to diethylthiophosphate (DETP) and trichloropyridinol with low/no toxicity, utilizing DETP as energy source for their growth (Fig 5).

**Fig. 2.**
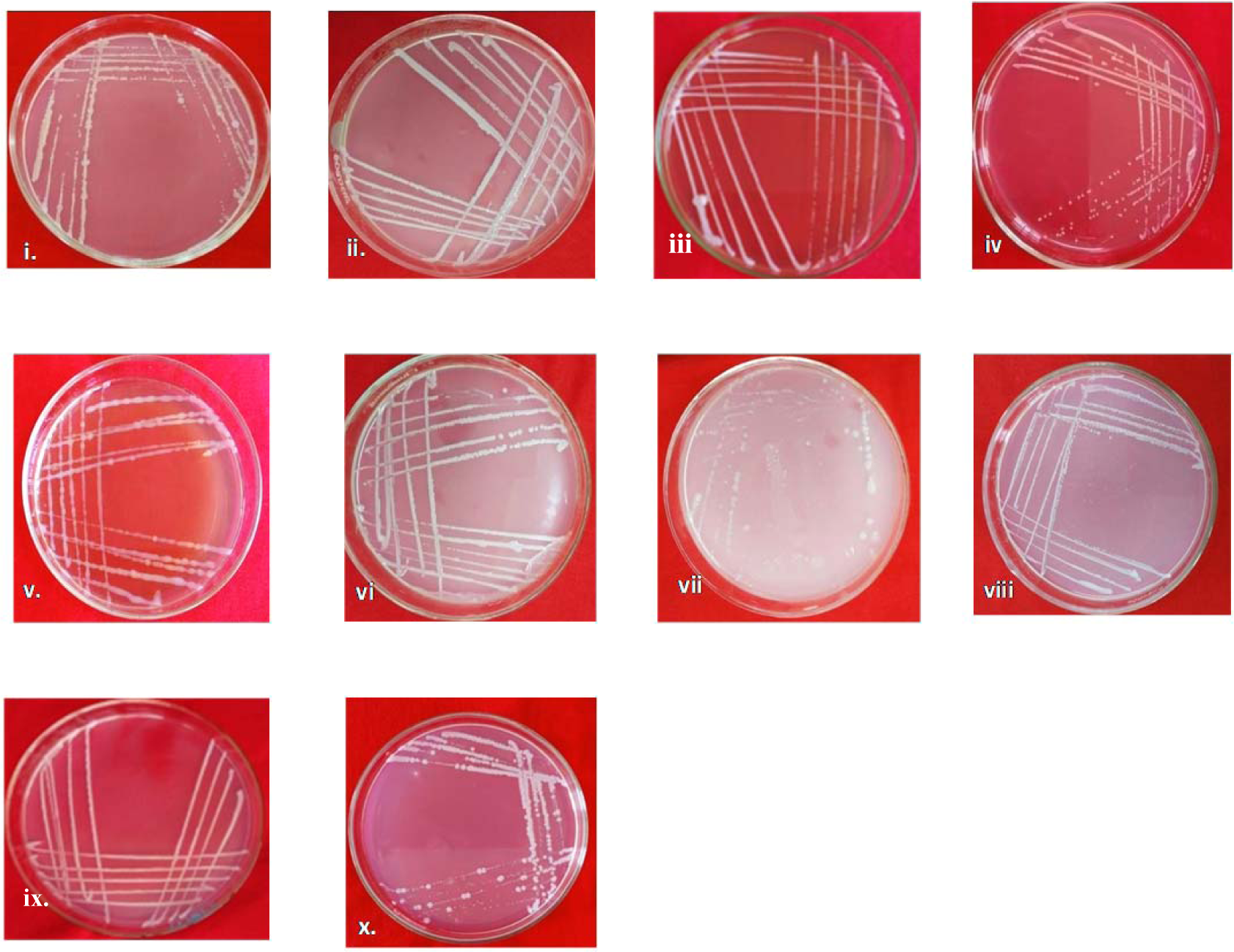
Growth of pure culture of OPDB isolates in chlorpyriphos amended (25 ppm) MS-medium source (i) TZ1 (ii) TZ2 (iii) TZ3 (iv) JHT1 (v) JHT2 (vi) JHT3 (vii) ICR1 (viii) ICR2 (ix) MG1, and (x) MG2. The annotations, i, ii, iii, and iv represent response at 100,300,500, and 700 ppm of chlorpyriphos, respectively.

**Fig. 3.**
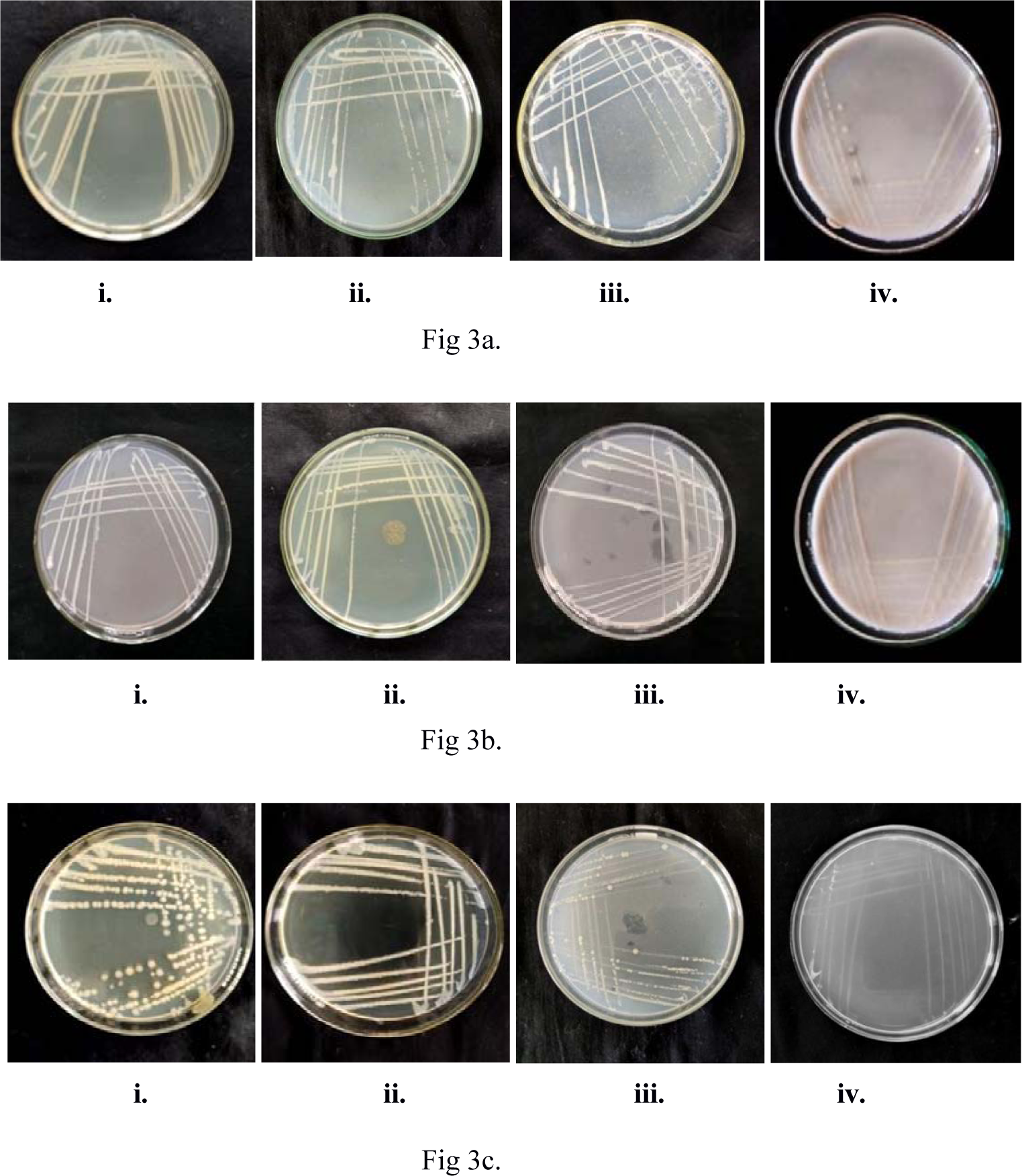
In-vitro growth of three selected OPDB isolates (JHT1, MG2, and TZ2) at four different concentrations of OP (100-700 ppm chlorpyrifos) in MS-medium. Fig 3a. *In vitro* growth Response of OPDB isolate (JHT1); Fig 3b. *In vitro* growth response of OPDB isolate (MG2); and Fig 3c. *In vitro* growth response of OPDB isolate (TZ2). The growth of OPDB isolate, TZ2 is significantly curtailed at 700 ppm of chlorpyriphos.

**Fig. 4.**
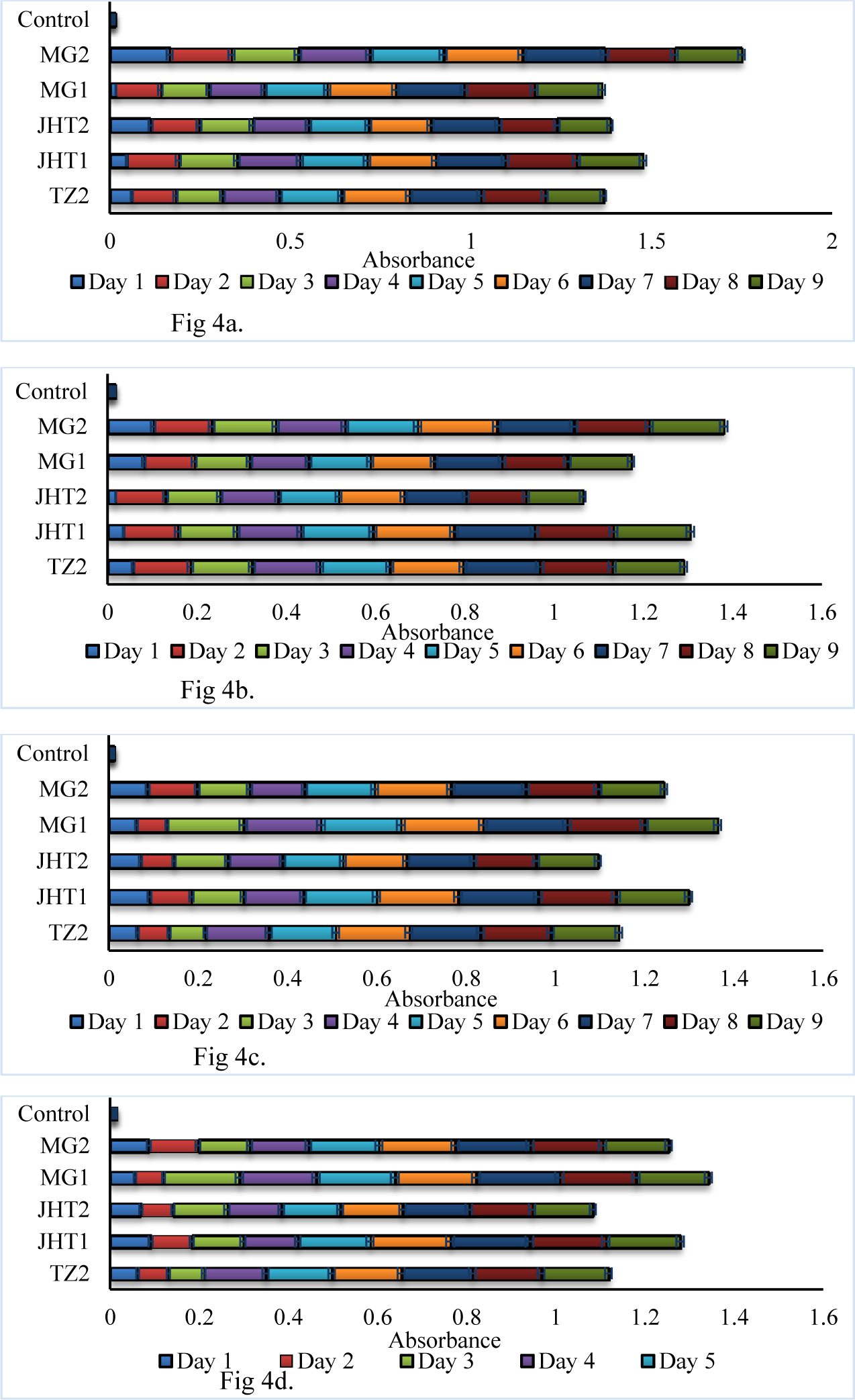
Growth response (absorbance as optical density calibrated against cfu/ml) of five efficient OPDB isolates at four different concentrations of chlorpyrifos (100,300,500, and 700 ppm) during 10-days of incubation study. Two OPDB isolates (JHT1 and MG2) displayed consistent growth in their colony population compared to other three isolates (MG1, JHT2, and TZ2) at all four concentrations of chlorpyrifos Fig 4a. Growth kinetics of the efficient OPDBs at 100 ppm of chlorpyrifos; Fig 4b. Growth kinetics of the efficient OPDBs at 300 ppm; Fig 4c. Growth kinetics of the efficient OPDBs at 500 ppm; and Fig 4d. Growth kinetics of the efficient OPDBs at 700 ppm.

**Fig.5.**
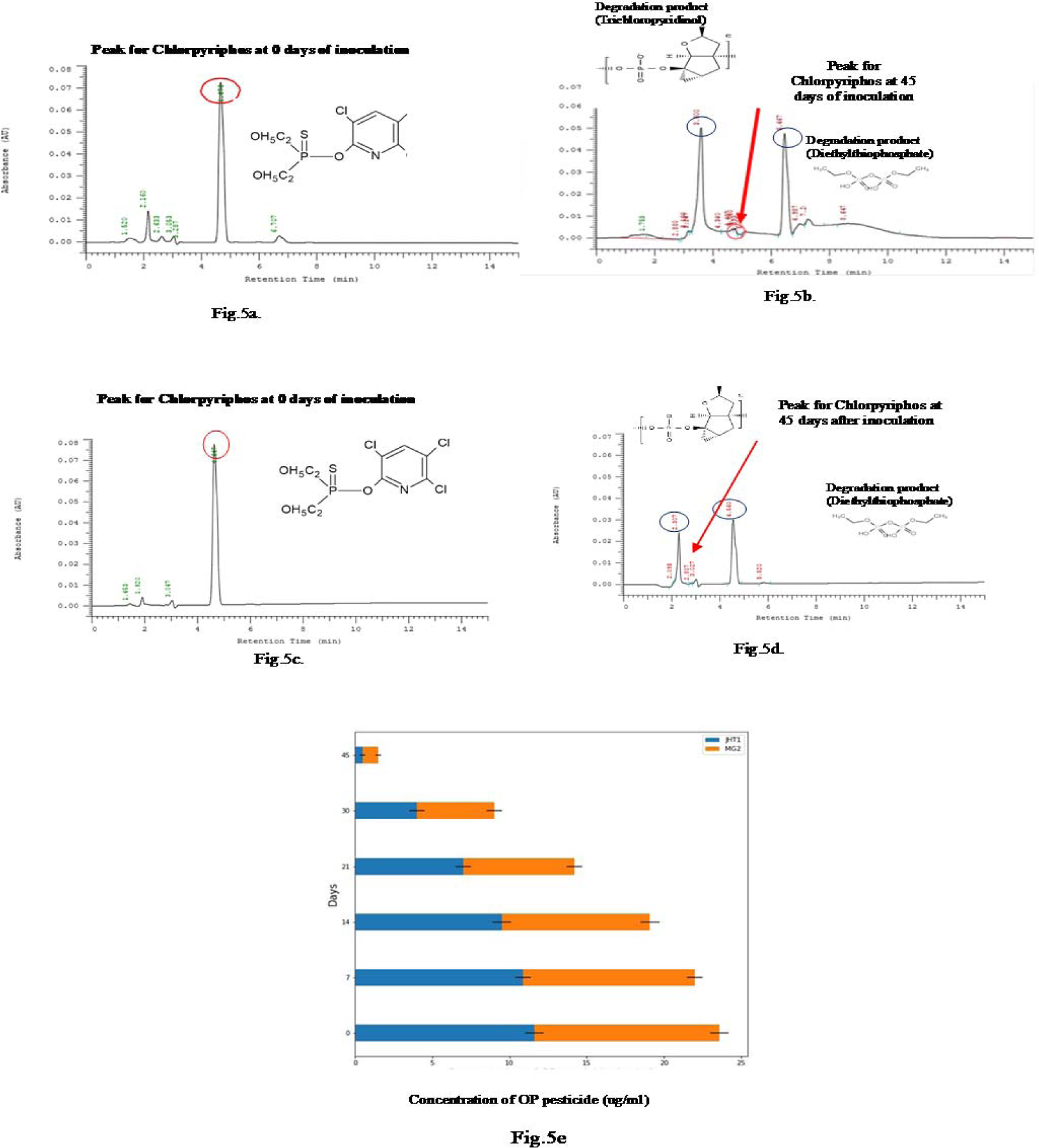
(5a-5e). Pesticide bio-degradation efficiency of two most efficient OPDB isolates (JHT1 and MG2) in 45-days of inoculation treatment under *in vitro* conditions. Fig.5a. Microbial degradation of pesticide by isolate JHT1at 0-day of inoculation. Presence of one high peak represents the dominance of chlorpyriphos pesticide. Fig.5b. Microbial degradation of pesticide by isolate JHT1at 45-days of inoculation. Presence of two lower peaks mark the dominance intermediate non toxic metabolites trichloropyridinol and diethylthiophosphate. Fig.5c. Microbial degradation of pesticide by isolateMG2 at 0 day of inoculation. Presence of one high peak represents the dominance of chlorpyriphos pesticide. Fig.5d. Microbial degradation of pesticide by isolate MG2 at 45-days of inoculation. Presence of two lower peaks demote the dominance of non toxic metabolites trichloropyridinol and diethythiophosphate. Fig.5e. Pesticide bio-degradation efficiency of two most efficient OPDB isolates (JHT1 and MG2) using HPLC-based chromatograms and efficiency progression during 45-days of inoculation.

### Characterization of screened OPDB

Two OPDB isolates (JHT1 and MG2) were further subjected to morphological, cultural, and molecular characterization (Fig. 6, 6a-6b) (Supplementary Table 3 and 4). Morphologically, both the isolates were observed Gram negative and rod-shaped. While, cultural characterization of JHT1 (greyish white, circular convex, opaque smooth, without pigment production) and MG2 (pale yellowish green, convex, rough edges) showed sharp contrast difference in their features. The phylogenetic analysis of two OPDB (JHT1 and MG2) using partial 16S rRNA gene sequences corroborated highest homology (more than 99% similarity) with *Achromobacter marplatensis, Am* (Fig. 6, 6a) and *Pseudomonas azotoformans*, *Pa* (Fig.6, 6b), carrying NCBI accession number of MW397524 and MW397525, respectively.

**Fig. 6.**
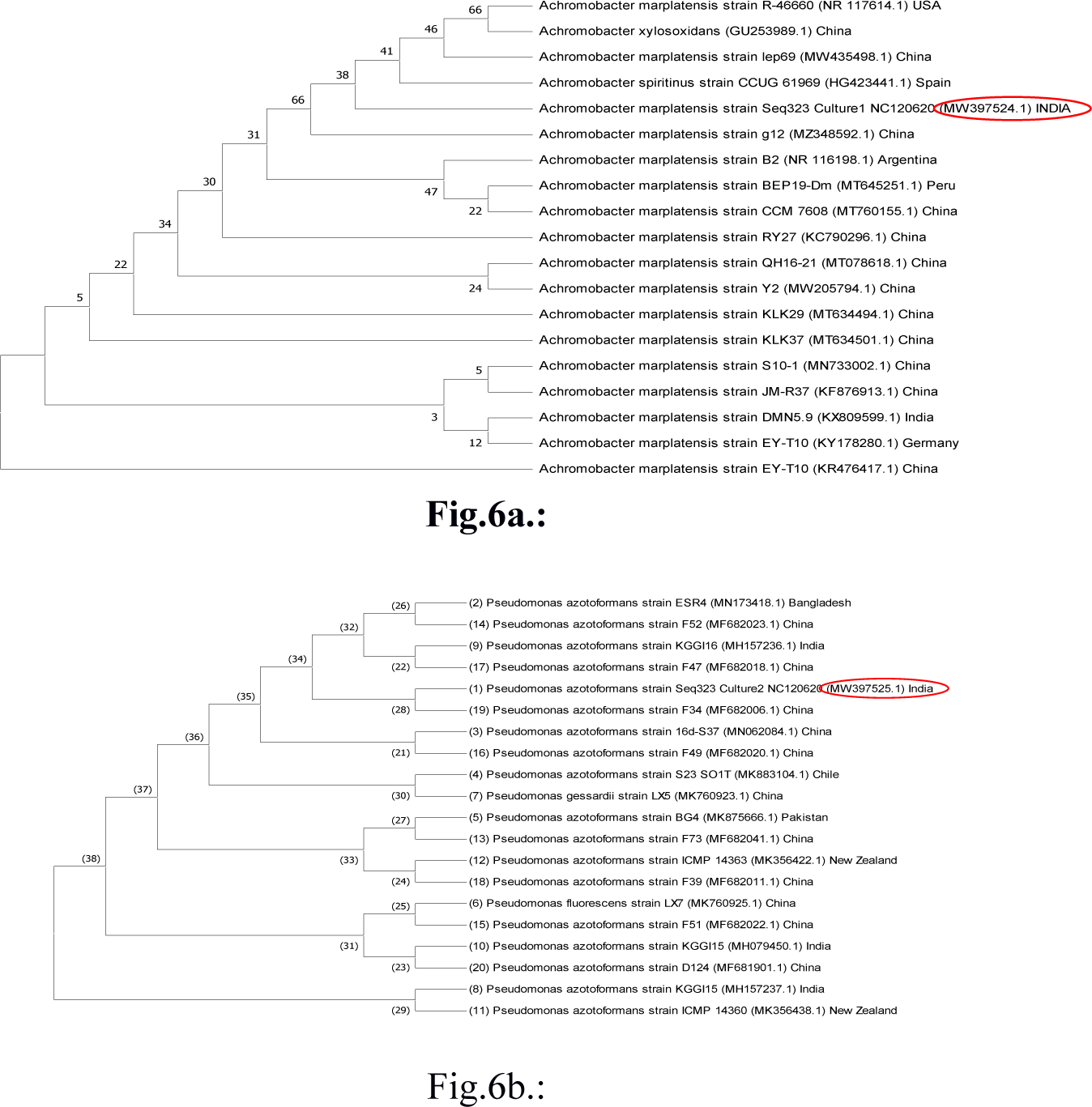
(6a-6b) Phylogenetic analysis of two most efficient OPDB isolates (JHT1, MW 397524 and MG2, MW 397525). Fig.6a. Phylogenetic tree showing genetic relationship of JHT1 isolate with other isolates using 500 bootstrap replicates; Fig.6b. Phylogenetic tree showing genetic relationship of MG2 isolate with other isolates using 500 bootstrap replicates

### MBCA-OPDB compatibility and their agronomic response

The compatibility between two MBCA*Pf*(KT258013) and *Th*(ON364138)] was established *in vitro* (Fig. 7a), which possess proven record of antagonism against *Rs*. Simultaneously, the compatibility between two most efficient OPDB (Fig. 7b), *Pf-Am-Pa* (Fig. 7c), *Th-Am-Pa* (Fig.7d), and *Pf-Th-Am-Pa* (Fig. 7e) were observed *in vitro*. These *in vitro* studies offered a natural choice for developing a suitable combination of MBCA and OPDB, evident from absence of inhibition zone around the microbial disk (Fig. 7, 7a-7e).

**Fig.7.**
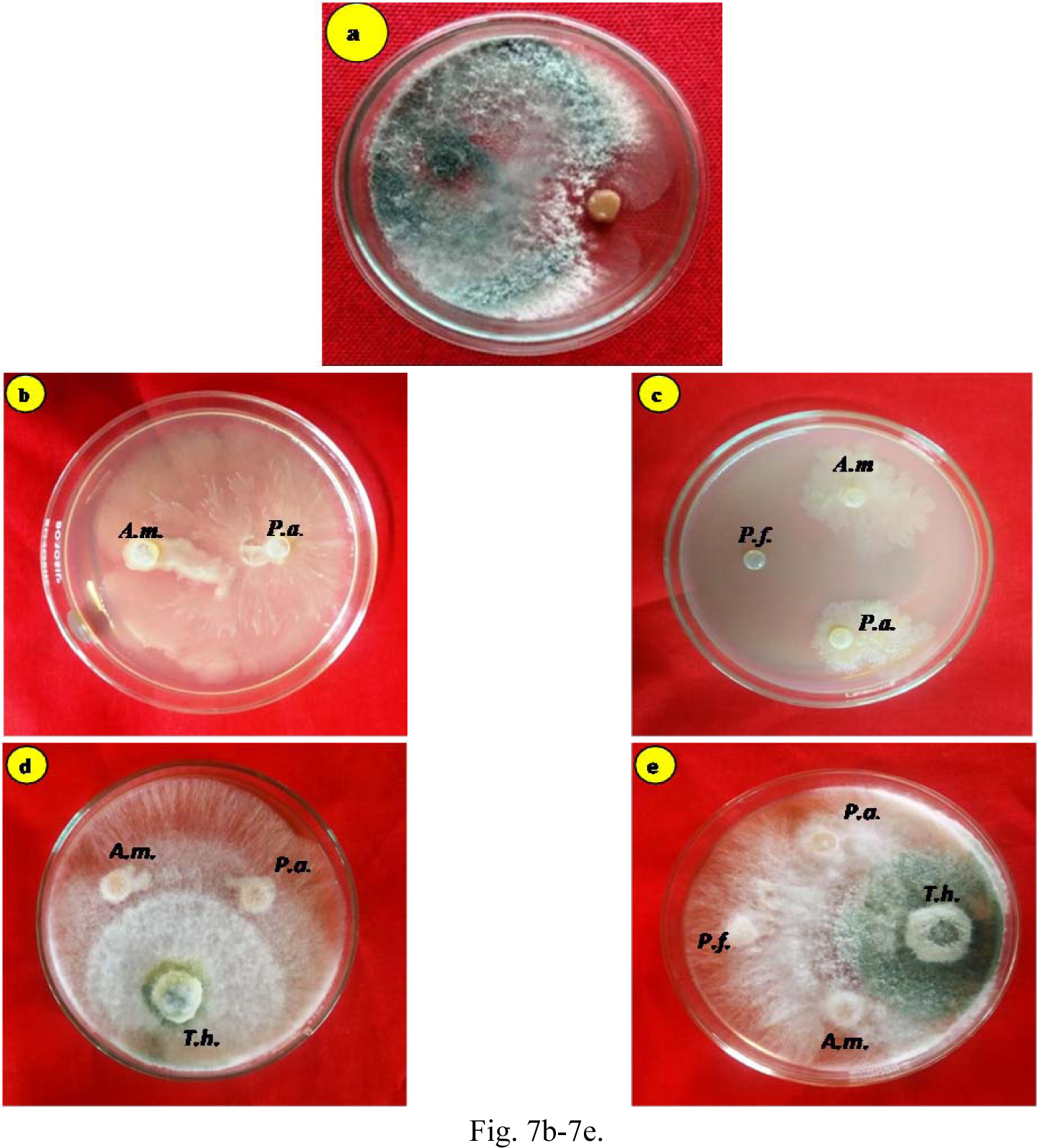
*In vitro* evaluation of compatibility between MBCA and OPDB isolates. Fig.7a. *In vitro* compatibility of Th (*Trichodermaharzianum)* and Pf (*Pseudomonas fluorescens*); Fig. 7b-7e. *In vitro* relationship between OPDBs isolates (Am, *Achromobacter marplatensis* and Pa, *Pseudomonas azotoformans*) and Am-Pa with MBCA (7c. Am-Pa-Pf: 7d. Am-Pa-Th and 7e. Am-Pa-Pf-Th). The overlapping in growth of colonies of Am×Pa×Pf×Th is clearly visible, offering a possible combination of OPDB and MBCA.

In a potted experiment, different combinations of MBCA (*Th* and *Pf*) and OPDB (*Am* and *Pa*) showed varying response on PWI, being lowestwith combination of *Pf + Th + Am + Pa* (T_5_) followed by *Pf + Th + Pa* (T_4_). However, the magnitude of reduction in PWI was significantly higher with *Pf* +*Th* (T_2_) compared to either *Pf +Th + Am* (T_3_) *or Pf + Th + Pa* (T_4_), displaying the coordinated response of MBCA with OPDB. The reduction in PWI was associated with an enhanced yield of brinjal and all the treated combination of *Pf + Th + Am + Pa* (T_5_) was most effective in enhancing the yield of brinjal followed by a combination of *Pf + Th + Pa* (T_4_). The other two combinations, *viz., Pf + Th* and *Pf + Th + Am* were however observed displaying higher magnitude of response over control treatment only. These observations followed the similar pattern of response with regard to yield attributing parameters, such as root and shoot dry weight, suggesting the superior agronomic response along with reduction in wilt disease of brinjal with combination of MBCA and OPDB over either of the two alone (Table 1).

**Table 1.** Response of PGPMs and OPDB on PWI, yield attributes, and yield of brinjal grown in OP-loaded potted soil

| Treatments | PWI<br>(%) | Yield attributing attributes |  | Yield<br>(kg/plant) |
| --- | --- | --- | --- | --- |
|  |  | Shoot dry<br>weight (g/plant) | Root dry weight<br>(g/plant ) |  |
| T<sub>1</sub>(Control) | 85.0e | 16.96e | 3.52e | 0.50de |
| T<sub>2</sub>(Pf-Th) | 15.0bc | 28.75d | 4.28c | 1.10bc |
| T<sub>3</sub>(Am- Pf-Th) | 20.0bcd | 27.95c | 4.07cd | 0.74d |
| T<sub>4</sub>(Pa- Pf-Th) | 10.0ab | 31.46b | 4.44b | 1.23b |
| T<sub>5</sub>(Am-Pa-Pf-Th) | 5.0a | 36.39a | 5.35a | 1.42a |
| S.Ed (+) | 4.49 | 0.32 | 0.13 | 0.16 |
| C.D. (P=0.05) | 9.86 | 0.67 | 0.27 | 0.33 |

*Pf*, *Pseudomonas flourescens* ; *Th* , *Trichoderma harzianum* ; *Am*, *Achromobacter marplatensis* ; *Pa* , *Pseudomonas azotoformans* ; and PWI , Percent wilt incidence . Combination of *Am* + *Pa* as OP-dregrading bacteria ( OPDB) isolated form OP-contaminated brinjal fields and *Pf* + *Th* as microbial biocontrol agents ( MBCA) out-performed rest of the other combinations, including either OPDB or MBCA alone. Means are separated using different alphabets through Duncans' Multiple Range Test

Different combinations of MBCA and OPDB were observed equally effective in degrading OP–pesticide to varying proportions. The combination carrying *Pf + Th + Am + Pa* (T_5_) was observed most efficacious in reducing the pesticide concentration to only 28.96% in 45 days of treatment (Fig. 8). Amongst the other treatment combinations, *Am* with *Pf + Th* (T_3_) showed greater efficacy than *Pa* with *Pf + Th* (T_4_) in reducing the concentration of OP-pesticide to 33.45% and 41.54%, respectively, in 45-days of treatment response. Our observations, hence, suggested combination of MBCA and OPDB in cleansing OP-pesticide from contaminated soil, thereby, facilitating abling environment for pathogenicity of wilt pathogen weakened through MBCA to ensure growth and yield

**Fig.8.**
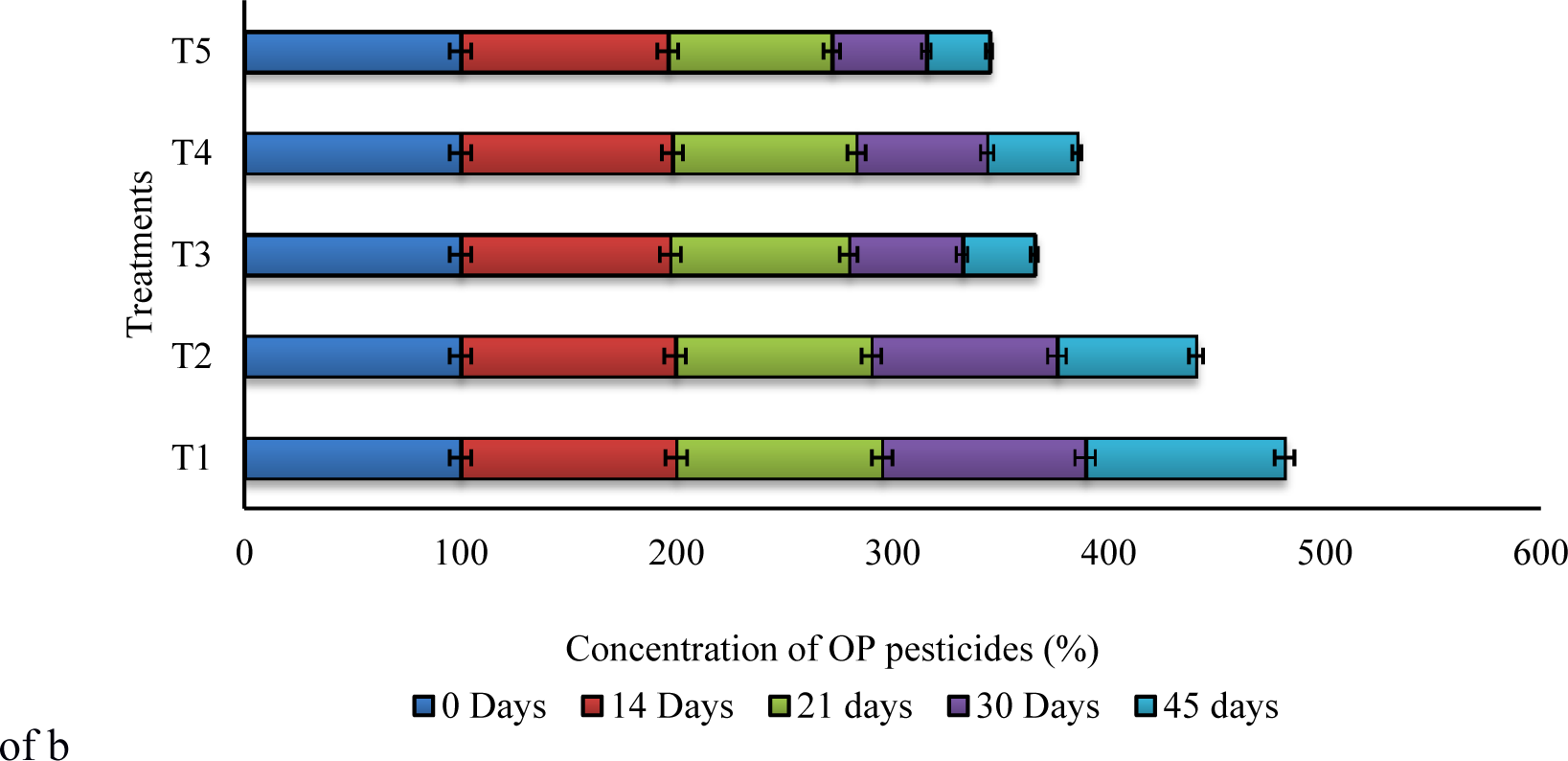
Effect of different combinations of MBCA (*Pf* and *Th*) and OPDB (*Am* and *Pa*) in biodegradation of OP. Treatments T_1_ involving combination of MBCA and OPDB significantly outplayed rest of the other treatments, viz., T_2_, T_3_ or T_4_ or T_5_ in terms of bio-efficiency. Treatments comprise of (T_1_- Control; T_2_- *Pf* + *Th*; T_3_- *Am*+ *Pf* + *Th;* T_4_- *Pa* + *Pf* + *Th;* T_5_- *Am* + *Pa* + *Pf* + *Th*). *Pf, Th, Am,* and *Pa* represent *Pseudomonas fluorescens, Trichoderma harzianum, Achromobacter marplatensis*, and *P. azotoformans*, respectively.

## DISCUSSION

OP-pesticides are reported highly effective against a wide range of insect pests, making them a popular choice in vegetable crops (10) and simultaneously causing multi-pronged health hazards to different agro-ecosystems through inhibition in acetylcholinesterase activity (41). Despite OP-residues known for relatively low persistence due to their water solubility, 70% of initially applied chlorpyriphos remained in soil after 18-months of application, with half-life of 116-1576 days (42), thereby, rendering them more vulnerable to human access (43,44). The bacterial wilt of brinjal crop caused by *Rs* pathogen is one of the serious endemic diseases, the causal agent is reported soil-borne in nature (14,26) with an extended survivability under favourable environmental conditions (18). According to Nahar *et al.* (45), improved nursery management practices(seeds treated with hot water and transplanted in nursery soil treated with *Th*) alone reduced bacterial wilt by 14–20 %, increasing marketable yield by 8–19 tons/ ha, and income by €1800-3700/ ha compared to farmers’ conventional practices (spraying of carbendazim). Such practices encouraged thepersistence of *Th* throughout the growing season and reduced the population of *Rs*, implying that the next crop would be raised at a much lower disease load.

The first successful attempt to degrade OP-compounds microbially was made in 1973 with the involvement of *Flavobacterium* sp., which laid the foundation of microbe assisted OP-degradation in liquid and soil cultures. The biochemistry of bacterial degradation of OP is more or less quite similar, where structurally similar enzyme called organophosphate hydrolase or phosphotriesterase catalyzes the first step of degradation (46). Persistence of OP –residues in 12-93% samples of brinjal and other vegetable growing soils with residues ranging from 0.384 to 0.980 mg/kg (0.05 mg/kg as maximum tolerance limit) is reported worldwide (47,48,49,50). Isolation of ten OPDB from OP-contaminated brinjal growing soils indicated the abundance of microbes carrying specific function of microbial degradation of OP-related residues. Previous studies reporting successful isolation of OPDB from varied soil environments with repeated pesticides application, lending strong support that pesticides are naturally degraded in any soil environment involving number of processes, *viz*., hydrolysis, oxidation, alkylation, and dealkylation (3,15) and extracellular / intracellular organo-phosphorus phosphatase enzymes, *viz.*, hydrolase, phosphotriesterase, carboxyl esterase, and phosphatase (51,35). Melghani *et al.* (51) established the bioremediation response of *Pa*in contaminated soil treated with 200 µg/g, removing 86.81% profenofos within 48-hrs of incubation through hydrolysis, including the mineralization of one of the metabolites like 4 bromo-2-cholorophenol in a MS-medium, supporting strong pesticide degrading ability of *Pa*in contaminated soil.

The isolated OPDB strains subjected to *in vitro* bioassay showing potential colony growth up to 700 ppm of chlorpyriphos were considered highly efficient in OP-biodegradation. Other studies with bacterial isolates growing upto700 ppm (47) and 1000 ppm concentration of chlorpyriphos (IRLM 1 and IRLM 3) in MS-medium (29) were reported efficient in degrading OP-pesticides. We further established the compatibility between two isolates identified as JHT1and MG2, however without any compatibility of MG2 with TZ2 growing upto 700 ppm. Such an observation provide further clues about mutually positive relationship between OP-degrading microbes as major players participating in microbial disintegration of pesticides. For example, the compatibility between four OPDB isolates viz., *Pseudomonas putid*a (NII 1117)*, Klebsiella* sp.(NII 1118)*., Pseudomonas stutzeri* (NII 1119), and *Pa* (NII 1120) with elevated organophosphorus hydrolase activity (0.171 units/mL/min.) was previously reported for effective OP-degradation in the presence of metabolites like chlorpyriphos-oxon and diethylphosphorothioate (52). Our two identified isolates registered 75-80% higher OP-degradation over control in 45-days, evident through reduced concentration of chlorpyriphos residues and various degradation metabolites. These results are quite comparable with some of earlier studies. The isolates hydrolyzing chlorpyriphos to diethylthiophosphate (DETP) and trichloropyridinol (12) and newly identified intermediate, 2,6-dihydroxy pyridine (32) during chlorpyriphos degradation with low/no toxicity, utilized these intermediates as carbon and phosphorus sources for their multiplication. In another study by Ifediegwu *et al.* (52) evaluating three pesticide degrading bacteria viz., *Pa, Serretia marcescens* (*Sm*), and *Klebsiella oxytoca* (*Ko*) in MS-medium supplemented with 20mg/L chlorpyriphos, which showed *Pa* exhibiting maximum growth in 10-days (degrading 60% of applied pesticide) compared to *Sm* and *Ko* with maximum growth after 6-days of incubation (degrading 53-54% of applied pesticide) in 14-days of incubation study. Following the efficacy of two isolates (JHT1 and MG2), their molecular characterization identified them as *Am* (MW397524) and *Pa* (MW397525), respectively. Some of the earlier studies (53, 54) characterized OPDB as *Am and Pa* using 16S rRNA with more than 99% homology expression.

Identified OPDB isolates were further observed having compatibility with MBCA and within MBCA (*Pf* versus *Th*). The compatibility of *Pf* and *Th*is widely reported by several workers (55, 56), revealing their dual functional ability, growth promotion and effectiveness against a variety of plant diseases. Yang *et al.* (55) examining the role of strain DSP3 (*Alcaligenes faecalis* treated at 10^8^ cfu/g) in the degradation of chlorpyriphos and 3,5,6-trichloro-2-pyridinol (TCP) under different culture and soil conditions, showed higher rate of pesticide degradation over untreated soil. The study further revealed that all tested OP-pesticides possessed diethyl phosphorothionate side chains with phosphotriester bond in all compounds except TCP, thus suggesting the major hydrolysis at phosphotriester bond. The combination of MBCA and OPDB evaluated for biodegradation of chlorpyriphos in soils of pot grown brinjal plants showed maximum efficacy with *Am* + *Pa* + *Pf* + *Th* followed by *Am* + *Pf* + *Th* and *Pa* + *Pf* + *Th* in decreasing order, however all displaying significantly superior response over control treatment. OP-pesticides usually degenerate by hydrolysis upon exposure to sunlight and air (3), in addition to natural degradation in residues of phosphomidon upto 97% in the foliage and silique of mustard crop at 15-days after spraying (57). The other studies on response of combination of four bacterial strains represented by *Pseudomonas* and *Achromobacter* genera showed degradation of 80% endosulfan (58) and 93-100% chlorpyrifos treated at the rate of 200-500 ppm within 42-days of application (10). These strains in our study also displayed a substantial plant growth promoting traits such as phosphate solubilization, indole acetic acid production, and ammonia production, both in absence as well as in the presence of chlorpyriphos on account of their cumulative effect of MBCA and OPDB (Supplementary Table 3 and Supplementary Table 4).

Amidst these limited studies, our studies evaluating combination of MBCA and OPDB against wilt disease of brinjal and decontamination of OP-residues showed lowest PWI translating into maximum agronomic response, put forth an option to develop microbial consortium loaded with traits of two tier functions. The antagonists, *Th* and *Pf* suppressing *Rs* are extensively reported via production of diffusible substances, overcrowding of pathogen, secreting lytic enzymes (β1,3-glucanases in lysis of pathogen cell wall), antibiotics, and toxic metabolites like cyanide (59,60,61). In our study, combination of *Pf* and *Th* expanded the functional corridor of these two antagonists via ammonia production, phosphate solubilization, and indole acetic acid production, either in absence or in the presence of chlorpyriphos, thereby, facilitated in optimizing the performance of host crop. Some of our recent studies have strongly advocated the application of microbesin a consortium mode proved far more efficacious (62, 63), since brinjal plants bioprimed with *Trichoderma* sp. and challenged with *Sclerotinia sclerotiorum* were observed having higher amount of shikimic acid coupled with defense-related enzymes *viz*., polyphenol oxidase, peroxidase, and phenylalanine ammonia lyase (60). Such combination of MBCA and OPDB are also likely to accelerate many of the fundamental plant physiological processes such as photosynthesis rate, stomatal conductance, transpiration, internal CO_2_concentration, water-use-efficiency, and nutrient uptake [70], besides solubilization of several plant nutrients, sequestration of iron through siderophore production, and growth hormones production (64, 65) pivotal in agronomic crop response. In a recent study, Das *et al.* (65) reported consortium of endophytes represented by *Acinetobacter*, *Enterobacter*, and *Klebsiella* genera, supplemented with 40% urea-treated eggplants displayed crop response similar to 100% urea-treated eggplants, thereby, saving 60% N-fertilizer, in addition to consuming 100% chlorpyrifos (50 mg/5 mL) within 14-days of exposure. Such observations hold a much greater promise towards commercial development of MBCA-OPDB based bioformulations with a cost effective delivery mechanism to ensure development of residue free vegetable crop production system.

## MATERIALS AND METHODS

### Bacterial wilt pathogen

Bacterial wilt pathogen, *Rs* was collected from Culture Bank of Department of Plant Pathology, Assam Agricultural University, Jorhat, India. Pathogenicity test of identified pathogen, *Rs* was carried out with one-month-old potted brinjal plants (var. Navkiran, wilt susceptible variety) carrying soil (Entisol) characterized by pH 6.4, sand 60.1%, silt 22.3%, clay 17.6%, and organic carbon 1.6% following root inoculation technique (30). Two sets, each having five plants were inoculated with suspension (12 x 10^7^cfu/mL prepared by adjusting the optical density of 0.50 at 425 nmof pathogenic bacterium, (*Rs*) by cutting the lateral roots and dipping the cut roots in 10 mL of bacterial suspension for 1-hr, alongside another set with sterile distilled water as standard check and kept under observations for development of symptoms in 15-days of incubation study. Re-isolation of pathogen was made from plants displaying wilt symptoms as per Koch’s postulation (31). The pure culture of *Rs* was undertaken in triphenyl tetrazolium chloride (TTC) slants and incubated for 48-hrs at 28±1^0^C with periodic subculturing to keep the pathogen in active form. Morphological, cultural, and biochemical characterization of *Rs* were carried out following Bergey’s Manual of Determinative Bacteriology (32) [Supplementary Table 1].

### Isolation of OPDB strains and in vitro screening for OP-degradation

The OPDB were isolated from ten brinjal fields, represented by three districts of Assamviz., Jorhat(26.72/26.80^0^,00/97’N;94.12/94.22^0^00/88’E), Darrang(26.52/26.53^0^,00/58 ‘N;92.13/92.16^0^ 00/22’E), and Sonitpur (26.44/26.63^0^, 12/45’N; 92.33/92.24^0^00/47’E) having history of OP-usage for five years. These sites were taxonomically represented by Entisol and Inceptisol soil orders as per USDA Soil Taxonomy. Five top soil samples (0-20 cm depth) collected from these fields and packed in polythene bags with ice pack were brought to laboratory, Department of Plant Pathology for further analysis. The collected soil samples varied in soil pH from 6.2-6.5, sand 62.1-68.6%, silt 20.1-23.2%, clay 15.8-17.2%, and organic carbon 1.5-1.8%, with predominantly sandy loam texture. The soil samples collected from different locations were mixed thoroughly, filled into glass vials (dimension: 2.5 cm diameter x 8 cm length)assoil columns (capacity, 50 g soil) enriched with native microflora, with weekly addition of synthetic solution of commercial grade OP pesticide (TRICEL a.i 20ECchlorpyrifos, 25 ppm) for 15-days as per the protocol described by Latifi et al. (33). The OPDB were isolated from these chlorpyrifos enriched soils through serial dilution. One mL from last dilution (10^8^) of enriched soil was aseptically transferred into culture plates with MS agar medium (34, 12), supplemented with chlorpyriphos (25 ppm) as the sole carbon source and incubated at 27±2°C for 24-hrs. The bacterial colonies growing on plates were purified in NA-medium and differentiated based on morpho-cultural characters.

The bacterial isolates displaying growth at 25 ppm concentration of chlorpyriphos in MS-medium were screened for their growth ability at different higher concentrations of pesticide (50 ppm, 100 ppm, 300 ppm, 500 ppm, 700 ppm, and 1000 ppm). The OPDB isolates exhibiting growth above 500 ppm were carried forward to test their growth kinetics spectrophotometrically (Model: Double Beam Spectrophotometer 2203, Systronics, India) in MS-broth at 50, 100, 300, and 500 ppm concentration of pesticides (35). The broth was aseptically inoculated with selected strains and incubated on orbital shaker at 150 rpm and 37°C. Two most efficient isolates exhibiting maximum growth by assimilating chlorpyriphos as sole carbon source under liquid MS-medium were selected for further analysis through High Pressure Liquid Chromatography (HPLC, Model: HITACHI Chromaster, Japan). The chlorpyriphos degrading ability of the most efficient OPDB-isolates was further confirmed through HPLC-based analysis as per protocol suggested by (36). Identification of chlorpyriphos degradation products was made based on retention time via Gas Chromatography-Mass Spectrometry (GCMS).

### Characterization of efficient OPDB

Selected OPDB-isolates were characterized through series of biochemical (Supplementary Table 2) and physiological tests (Supplementary Table 3) using Bergey’s Manual of Systematic Bacteriology (37,38). While, molecular identification was performed using 16S rRNA amplification after extracting genomic DNA through Bacterial Genomic DNA Extraction Kit (QiaGen^TM^ Bacterial DNA Isolation Kit) and purified using a DNA Purification Kit (GeneiPure^TM^ Bacterial DNA Purification Kit). The sequence of the isolates submitted to Gene Bank to search similar sequences using the BLAST algorithm1 and was compared with known sequences of NCBI Gene Bank database (33). The top hit known sequences of BLAST^®^ search results for each strain were further multiple aligned using CLUSTAL software and constructed phylogenetic relationship using MEGA 7 Windows version software through neighbor-joining procedure.

### Combination of OPDB with MBCA

The efficient OPDBs were tested for their compatibility with each other as well the MBCA viz., *Th* (NCBI accession no ON364138) and *Pf* (NCBI accession noKT258013) procured from the author’s laboratory (Biocontrol Laboratory), Department of Plant Pathology, AAU, Jorhat (India) having established antagonism against *Rs* (37) were tested *in vitro* for their compatibility with two selected OPDB (OPDB1 and OPDB2) through dual culture technique (34) using nutrient agar (NA) and Potato Dextrose Agar (PDA) as basal media, placing the OPDB and MBCA beads from NA and PDA culture plates (stored at 27°C for 1 day and 25°C for 5-days, respectively) opposite to each other,1.5 cm from the edge of the culture plate. The compatibility was determined through development of inhibition zone after incubation at 25°C for 5 days. A total of 4 treatments comprising i. Control, ii. OPDB1+ OPDB2, iii. *Pf*+ OPDB1 +OPDB2, iv. *Th*+ OPDB1 + OPDB2, and v. *Pf*+ *Th*+ OPDB1 + OPDB2, replicated 5 times were tested through completely randomised design (CRD). The efficacy of these treatments was worked out based on colony growth following 120-hrs of incubation.

Pure culture of MBCA and OPDB was maintained in NA-slants and incubated at 28±1°C for 24 hrs with periodical addition of sterile distilled water to maintain an un-interrupted colony growth and prepare bioinoculants using the protocol suggested by (34) with some modifications. The suspension of bacterial inoculants (OPDB), and bioagent (*Pf*) each amounting 15 mL were aseptically added to 1L of NA-broth and King’s broth, respectively, and incubated at 28±1°C for 72-hrs to obtain the population of 1 × 10^7^colony forming units (cfu/mL), while PD-broth was used for *Th*. Each of *Pf*, *Th,* OPDB1, and OPDB2 (10 mL each in 1:1:1:1 ratio)along with1% carboxy-methyl cellulose as sticker and2% mannitol as osmoticant were aseptically inoculated in the substrate containing PD-broth maintained at 28±1°C for 7-days to develop the consortia combination as per procedural steps outlined by Sharma *et al.* (39). The shelf life of consortia combination was also assessed (Supplementary Table 4).

### Evaluation of combined MBCA - OPDB against Rs and OP-degradation

The compatible combinations of MBCA and OPDB thus developed were tested against bacterial wilt of brinjal and degradation of OP through pot (dimension: 45cm length x 25 cm diameter) grown brinjal (wilt susceptible variety, Navkiran) under net-house conditions (optimum temperature 27-30°C, photoperiod 7.2 hrs, and relative humidity 70-80%). The pot soil (Entisol, pH 6.5, 58.4% sand, 24.8% silt, 16.8% clay, and 1.8% organic carbon) was autoclaved at 121°C and 15 psi pressure for two consecutive days. A total of 5 treatments consisting of: T_1_(Control), T_2_ (*Pf* + *Th*), T_3_ (*Pf* + *Th +* OPDB1), T_4_ (*Pf + Th+* OPDB2), and T_5_ (*Pf + Th +* OPDB1 + OPDB2), replicated five times, were tested through CRD. Seven-days-old seedlings were inoculated with *Rs* (1x 10^8^cfu/mL) by root clip method (40) and transplanted into sterilized pot (UV sterilized for 30 minutes for 3 consecutive days) containing sterilized soil treated with 600 ppm chlorpyriphos along with recommended fertilizers (20gN, 30gP_2_O_5_, 10gK_2_O, and 5g ZnSO_4_/seeding/pot). Bioinoculants were applied as seed treatment for 1-hr (ST, 20mL/100 seeds) and drying under shade for 2-hrs, seedling root treatment (RT, 2% solution at100mL/100seedlings) by root dip for 1-hr followed by shade drying for another 1-hr, and soil application (SA, 20 mL of 2% consortia inoculants) of inoculants applied at 20- and 40-days after transplanting.

Number of wilted plants was recorded and expressed as percent wilt incidence (PWI): No. of plants wilted/Total number of plants×100. Different growth attributing characters, viz., number of leaves, number of branches, shoot dry weight, root dry weight and yield were recorded at 60 days of maturity after transplanting. OP-biodegradation was assessed in soil filtrates through HPLC following the methodology as suggested by Latifi *et al.* (33). The population assay of bacterial wilt pathogen *Rs* was worked out at 0-day and 60-days after planting (Supplementary Table 7).

### Statistical analysis

Data generated were subjected to statistical analysis for computation of critical difference (F-test) using SAS software (v8.1, SAS Institute, North Carolina, USA). Duncan’s Multiple Range Test was performed to separate the means at 5% level of significance.

## CONCLUSION

Bio-scavenging of pesticide residues and mending such contaminated soil for optimised crop performance involve two contrasting issues to be addressed. A media enrichment method used to isolate OPDB from OP (chlorpyriphos)-contaminated soil to utilize OP-residues effectively aided in decontaminating the pesticide-polluted soil. The combined response of MBCA and OPDB provided some useful insights for both, microbial degradation OP-residues and reduction in incidence of bacterial wilt disease of brinjal. Such a study as a concept offers an exciting scope in creating a dynamic biological barrier against heavy use of pesticides on one hand and developing strong plant immune through MBCA as second line of safeguard for robust green technology-based vegetable production system. We can provide better dynamism to such two-pronged studies with development of nutrient fortified bioformulation, duly evaluated through long term field studies involving multiple cropping sequences.

## SUPPLEMENTAL MATERIALS

Data pertaining to supplemental materials are supplied through Supplementary Table 1-7.

### Authors contributions

The first author (SSA) having carried out this study as part of Ph.D under the supervision of second author (PB). Both the authors conceptualised the study, planned, and executed. Data generated out of the study were statistically analysed by both the authors (SSA and PB). Both the authors developed the manuscript and edited for final submission.

## ACKNOWLEDGEMENTS

Authors sincerely acknowledge Department of Plant Pathology, Assam agricultural University, Jorhat, Assam, India for providing all the necessary facilities during the course of the study.

## REFERENCES

1. Aktar W, Sengupta D, Chowdhury A. 2009. Impact of pesticides use in agriculture: their benefits and hazards. Interdiscip Toxicol 2(1): 1–12.

2. Nicolopoulou –Stamati P, Maipas S, Kotampasi C, Stamatis P, Hens L. 2016. Chemical pesticides and human health: The urgent need for a new concept in agriculture. Front Public Health 4: 148. doi 10.3389/fpubh.2016.00148.

3. Kumar S, Kaushik G, Dar M A, Nimesh S, Lopez-Chuken U J, Villarreal-Chiu J F. 2018. Microbial degradation of organophosphate pesticides: a review. Pedosphere 28(2): 190–208.

4. Ganesan K, Raza S K, Vijayaraghavan R. 2010. Chemical warfare agents. J Pharm BioAllied Sci 2: 166–178.

5. Jayashree R, Vasudevan N. 2007. Effect of tween 80 added to the soil on the degradation of endosulfan by *Pseudomonas aeruginosa*. Inter J Environ Sci Technol 4(2):203–210.

6. Watts M. 2021. Chlorpyrifos as a Possible Global POP. Pesticide Action Network North America, Oakland, CA, USA. www.pan-europe, 2021, pp. 1–34.

7. Pérez-Lucas G, Vela N, El Aatik A, Navarr S. 2018. Environmental risk of groundwater pollution by pesticide leaching through the soil profile. In: Pesticides Use and Misuse and their Impact in the Environment. Intech Open doi: 10.5772/intechopen.82418.

8. Boedeker W, Clausing M W P, Marquez E. 2020. The global distinction of acute unintentional pesticide poisoning estimations based on a systematic review. BMC Pub Health 20: 1875.doi.org/10.1186/s12889-020-09939-0.

9. Randhawa M A, Zaman M, Anjum F M, Ashgar A, Sazid M W. 2015. Organophosphate pesticide residues in okra and brinjal grown in peri-urban environment of big cities of Punjab. J Chem Soc Pak 37: 574–578.

10. Akbar S, Sultan S. 2016. Soil bacterial showing a potential of chlorpyrifos degradation and pant growth enhancement. Braz J Microbiol 47: 563–570.

11. Karami-Mohajeri S, Abdollahi M. 2011. Influence of organophosphate, carbamate, and organochlorine pesticides on cellular metabolism of lipids, proteins, and carbohydrates: a systematic review. Hunan Exp Toxicol 30: 1119–1140.

12. Singh K, Brajesh K, Walker A, Alum J, Morgan W, Wright D J. 2004. Biodegradation of chlorpyrifos by *Enterobacter* strain B-14 and its use in biodegradation of contaminated soils. Appl Environ Microbiol 70: 4855–4863.

13. Das G, Islam T. 2014. Relative efficacy of some newer insecticides on the mortality of jassid and white fly in brinjal. Intern Res J Biol Sci 4(3): 89–93.

14. Bashar M A, Chakma M. 2010. *In vitro* control of *Fusarium solani* and *F. oxysporum*, the causative agents of brinjal wilt. Dhaka Univ J Biol Sci 23: 53–60.

15. Chandrashekar, K N, Prasannakumar M K, Deepa M, Vani A, Khan A N A. 2012. Prevalence of races and biotypes of Ralstonia solanacearum in India. J Pl Prot Res 52:53–58.

16. Jamil Arshi, Manish Kumar. 2021. Suppression of Fusarium wilt of eggplant using *Trichoderma harzianum* and carbendazim. Intern. J Veg Sci doi 2021 .org./10.1080/19315260.2021.1920541

17. Singh N, Phukan T, Sharma P L, Kabyashree K, Barman A, Kumar R, Ray S K. 2018. An innovative root inoculation method to study *Ralstonia solanacearum* pathogenicity in tomato seedlings. Phytopathol 108(4): 436–442.

18. Sarkar S, Chaudhuri S. 2016. Bacterial wilt and its management. Curr Sci 110: 1439–1445.

19. Pradhanang P, Ji P, Momol, M, Olson M, Mayfield L J, Jones J. 2005. Application of acibenzolar-s-methyl enhances host resistance in tomato against *Ralstonia solanacearum*. Pl Dis 89: 989–993.

20. Chen S, Qi G, Ma G, Zhao X. 2020. Biochar amendment controlled bacterialwilt through changing soil chemical properties and microbial community. Microbiol Res 231: 126373.

21. Schreinemachers P, Chen P H, Loc N T T, Buntong B, Bouapao L, Guatam S, Nhu T L, Pinn T, Vilaysone P, Srinivasan R. 2017. Too much to handle? pesticide dependence of smallholder vegetable farmers in Southeast Asia. Sci Total Environ 593: 470–477.

22. Bora Popy, Bora L C, Deka P C, Bikram B, Sharma A K, Dutta H S, Debahaj B (2016b) Efficacy of *Pseudomonas fluorescens* and *Trichoderma viride* based bioformulation for management of bacterial wilt disease of ginger. Intern J Pl Sci (Muzaffarnagar) 11(2): 180–186.

23. Saddler G S. 2005. Management of bacterial wilt disease. In: Bacterial Wilt Disease and the Ralstonia solanacearum Species Complex (ed: Allen, C., Prior, P., Hayward, A. C.), American Phytopathological Society, APS Press, St. Paul, MN, 2005, pp. 121–132.

24. Lemessa F, Zeller W. 2007. Isolation and characterization of *Ralstonia solanacearum* strains from Solanaceae crops in Ethiopia. J Basic Microbiol 474:0–49.

25. Chakravarty G, Kalita M C. 2011. Comparative evaluation of organic formulations of *Pseudomonas fluorescens*based biopesticides and their application in the management of bacterial wilt of brinjal (*Solanum melongena* L.). African J Biotechnol 107: 174–7184.

26. Bora Popy, Bora L C, Begum M. 2013. Eco-friendly management of soil borne diseases in brinjal through application of antagonistic microbial population. J Biol Control 27(1): 29–34.

27. Singh S, Sinha A, Yadav S M, Singh B K, Singh H, Srivastava S, Kumar R. 2017. Antagonistic behavior of different bioagents against dominant seed borne fungi of mungbean seeds under in vitro condition. Proc Nat Acad Sci 87(2): 599–602.

28. . Papavizas G. 1985. *Trichoderma* and *Gliocladium*: biology, ecology, and potential for biocontrol. Ann Rev Phytopathol l23(1): 23–54.

29. Bora L C, Bora Popy, Gogoi M. 2020. Potential of *Trichoderma* spp. for pest management and plant growth promotion in NE India. In: Advances in Trichoderma research. (Ed: P. Sharma and Anil Sharma), Springer Publication, New York, USA, 2020, pp. 205–220.

30. Winstead N N, Kelman A. 1952. Inoculation technique for evaluating resistance to *R. solanacearum* (*Pseudomonas solanacearum*). Phytopathol 42: 628–634.

31. Allyson L, Byrd Julia, Segre A. 2016. Adapting Koch’s postulates. Science 351: 6270, 224–226 doi: 10.1126/science.aad6753.

32. Mali H, Shah C, Patel D H, Trivedi U, Subramanian R B. 2022. Degradation insight of organophosphate pesticide chlorpyrifos through novel intermediate by *Arthrobacter* sp. HMO1. Bioresour Bioprocess 9: 31. doi.org/10.1186/540643-022-00515-5.

33. Latifi A M, Khodi S, Mirzaei M, Miresmaeili M, Babavalian H. 2012. Isolation and characterization of five chlorpyrifos degrading bacteria. African J Biotechnol 11(13):3140–3146.

34. Karpouzas D G, Walker A. 2000. Factors influencing the ability of *Pseudomonas putida* strains epI and II to degrade the organophosphate ethoprophos. J Appl Microbiol 89(1): 40–48.

35. Yadav S, Verma S K, Chaudhary H S. 2015. Isolation and characterization of organophosphate pesticides degrading bacteria from contaminated agricultural soil. J Biol Sci 15(3): 113–118.

36. Li X, Jing J, Gu L, Ali S, He J, Li S. 2008. Diversity of chlorpyrifos degrading bacteria isolated from chlorpyrifos contaminated samples. Intern. Biodeterior. Biodegradation 62: 331–335.

37. Sneath P H A. 1986. Bacterial nomenclature. In: Bergeys Manual of systematic Bacteriology (Sneath, P.H.A., Mair, N.S., Sharpe, M.E., Holt, J.G., Ed), 1986, Vol 1, pp. 83–88.

38. Wayne L G, Kubica G P. 1986. Family Mycobacteriaceae. In: Bergeys Manual of systematic Bacteriology(Sneath, P.H.A., Mair, N.S., Sharpe, M.E., Holt, J.G., Ed), 1986, Vol 2, pp. 1435-1457.

39. Sharma P, Bora, L C, Puzari K C, Baruah A M, Baruah R, Talukdar K, Phukan A. 2017. Review on bacterial blight of rice caused by *Xanthomonas oryzae*pv. *oryzae*: different management approaches and role of *Pseudomonas fluorescens* as a potential biocontrol agent. Int J Curr Microbiol Appl Sci 6: 982–1005.

40. Bora Popy, Bora L C, Deka P C. 2016a. Efficacy of substrate based bioformulation of microbial antagonists in the management of bacterial disease of some solanaceous vegetables in Assam. J Biol Control 30(1): 49–54.

41. Sidhu G K, Singh S, Kumar V, Dhanjal D S, Datta S, Singh I. 2019. Toxicity, monitoring and biodegradation of organophosphate pesticides: a review. Crit Rev Environ Technol 49: 1135–1187.

42. Racke K D, Lubinski R N, Fontaine D D, Miller J R, McCall P J, Oliver G R. 1993. Comparative fate of chlopyrifos insecticide in urban and agricultural environments. ACS Symp Series 522: 70–85.

43. EPA (Enviornmental Protection Agency) Fate, transport and transformation, test guidelines. United States Environment Protecion Agency, 2008, 7101, USA.

44. Tse H, Comba M, Alaee M. 2004. Method for the determination of organophosphate insecticides in water, sediment and biota. Chemosphere 54: 41–47.

45. Nahar N, Islam Md R, Uddin M M, Jong P de, Struik P C, Stomph T J. 2019. Disease management in egg plant (*Solanum melongena* L.) nurseries also reduces wilt and fruit rot in subsequent plantings in Bangladesh. Bangladesh Crop Prot 120: 113–124.

46. Singh B K, Walker A. 2008. Microbial degradation of organophosphorus compounds. FEMS Microbiol Rev 30: 428–471.

47. Gowda AS R, Somashekar R K (2012) Evaluation of pesticide residues in farmgate samples of vegetable sin Karnatake, India. Bull Environ Contamin Toxicol 89: 626–632.

48. Prado-Lu J L D. 2014. Insecticide residues in soil, water and eggplant fruits and farmers health effects due to exposure to pesticides. Environ Health Preven Medicine 20:53–62.

49. Bhandari G, Zomer P, Atreya K, Mol H G, Yang X, Geissen V. 2019. Pesticide residues in Napalese vegetables and potential health risks. Environ Res 172: 511–521.

50. Habib M, Kaium A, Khan M S I, Prodhan M D H, Begum N, Chowdhury M T I, Isalm M A. 2021. Residue level and health risk assessment of organophosphorus residues in eggplant and cauliflower collected from Dhaka city of Bangladesh. Food Res 5: 369–377..

51. Melghani S, Chatterjee N, Hu X, Zejiao L. 2009. Isolation and characterization of a profenofos degrading bacterium. J Environ Sci 21: 1591–1597.

52. Ifediegwu M C, Agu K C, Wah N, Mbachu A E, Okeke C B, Anaukwu C, Uba P O, Ngenegbo U, Nwankwo O M. 2015. Isolation, growth and identification of chlorpyrifos degradating bacteria from agricultural soil in Anambra state, Brazil. Universal J Microbiol Res 3: 46–52.

53. Ma Y, Li E, Qi Z, Li H, Wei X, Lin W, Zhao R, Jiang, A, Yang H, Yin Z, Yuan J, Zhao X. 2016. Isolation and molecular characterization of *Achromobacter* phage phiAxp-3, an N4-loke bacteriophage. Sci. Rep. 6: 274–276.

54. Barua L, Bora D K. 2008. Comparative efficacy of *Trichoderma harzianum* and *Pseudomonas fluorescens* against *Meloidogyne incognita* and *Ralstonia solanacearum* complex in brinjal. Indian J Nematol 38: 86–89.

55. Yang L, Zhao Y H, Zhang B X, Zhang X. 2005. Isolation and characterization of a chlorpyrifos degrading bacteria and its bioremediation application in the soil. FEMS Micorbiol Letters 251: 67–73.

56. Yendo S, Ramesh, G C, Pandey, B R. 2018. Evaluation of *Trichoderma* spp., Pseudomonas fluorescens and Bacillus subtilis for biological control of Ralstonia wilt of tomato. F1000Research6:2028. https://doi.org/10.12688/f1000research.12448.3

57. Singh N S, Singh D K. 2011. Biodegradation of endosulfan and endosulfan sulphate by *Achromobacter xylosoxidans* strain C8B in broth medium. Biodegradation 22: 845–857.

58. Olanrewaju S O, Glick B R, Babalola O O (2017) Mechanism of action of plant growth promoting bacteria. World J Microbiol Biotechnol 33: 197–206.

59. Kohl J, Kolnaar R, Ravensberg W J (2019) Mode of action of microbial biological control agnets against plant diseases: Relevance beyond efficacy. Front Pl Sci 9:1801 doi:10.3389/fpls.2018.01801.

60. Bora Popy, Saikia K, Hazarika H. 2019. Ragesh, G. Exploring potential of bacterial endophytes in disease management of horticultural crops. Curr Hort 7: 32–37.

61. Bora Popy, Bora L C. 2021. Microbial antagonists and botanicals mediated disease management in tea, *Camellia sinensis* (L.) Kuntze.: an overview. Crop Prot 148: 105711.

62. Bora Popy, Bora L C, Bhuyan R P, Hashem A, Abd-Allah E F. 2022. Bioagent consortia assisted suppression in grey blight disease with enhanced leaf nutrients and biochemical properties of tea (*Camellia sinensis*). Biol Control 70:104907. https://doi.org/10.1016/j.biocontrol.2022.104907.

63. Singh S P, Keswani C, Singh S P, Sansinenea E, Hvat T X. 2021. *Trichoderma* spp mediated induction of systemic defense response in brinjal against Sclerotinia sclerotiorum.Curr Res Microbiol Sci 2:100051.doi:10.1016/jicrmicr.2021.100051.

64. Woo S, Pepe O. 2018. Microbial consortia: Promising probiotics as plant biostimulants for sustainable agriculture. Front Pl Sci 9,1801. doi:3389/fpls.2018.01801.

65. Das S R, Haque Md A, Akbor Md A, Abdullah Al-Mamun, Md Debnath C D, Hossain Md S, Hasan Z, Rahman A, Isalm Md A, Hossain Md Al-Amin, Yesmin S, Nahar M N E, Cho K M. 2022. Organophosphorus pesticides mineralising endophytes and rhizsopheric soil bacterial consortium influence eggplant growth promotion. Arch Microbiol 204: 199. doi.org/10.1007/s00203-022-02809-w.

